# Structural basis for activation and gating of IP_3_ receptors

**DOI:** 10.1101/2021.10.18.464806

**Authors:** Emily A. Schmitz, Hirohide Takahashi, Erkan Karakas

## Abstract

Calcium (Ca^2+^) is a universal and versatile cellular messenger used to regulate numerous cellular processes in response to external or internal stimuli. A pivotal component of the Ca^2+^ signaling toolbox in cells is the inositol 1,4,5-triphosphate (IP_3_) receptors (IP_3_Rs), which mediate Ca^2+^ release from the endoplasmic reticulum (ER), controlling cytoplasmic and organellar Ca^2+^ concentrations^1-3^. IP_3_Rs are activated by IP_3_ and Ca^2+^, inhibited by Ca^2+^ at high concentrations, and potentiated by ATP^1-3^. However, the underlying molecular mechanisms are unclear due to the lack of structures in the active conformation. Here we report cryo-electron microscopy (cryo-EM) structures of human type-3 IP_3_R in multiple gating conformations; IP_3_-ATP bound pre-active states with closed channels, IP_3_-ATP-Ca^2+^ bound active state with an open channel, and IP_3_-ATP-Ca^2+^ bound inactive state with a closed channel. The structures demonstrate how IP_3_-induced conformational changes prime the receptor for activation by Ca^2+^, how Ca^2+^ binding leads to channel opening, and how ATP modulates the activity, providing insights into the long-sought questions regarding the molecular mechanism of the receptor activation and gating.

## Introduction

IP_3_Rs are intracellular Ca^2+^ channels, predominantly localized to the ER and activated by the binding of IP_3_ generated in response to external stimulation of G-protein coupled receptors^1-3^. Opening of the IP_3_Rs results in the rapid release of Ca^2+^ from the ER into the cytoplasm triggering diverse signaling cascades to regulate physiological processes such as learning, fertilization, gene expression, and apoptosis. Dysfunctional IP_3_Rs cause abnormal Ca^2+^ signaling and are associated with many diseases, including diabetes, cancer, and neurological disorders^4,5^. There are three IP_3_R subtypes (IP_3_R-1, −2, and −3) that share 60-70% sequence identity, form homo- or hetero-tetramers, exhibit different spatial expression profiles, and are involved in different signaling pathways^1-3^. Each IP_3_R subunit is about 2,700 amino acids in length and contains a transmembrane domain (TMD) and a large cytoplasmic region comprising two β-trefold domains (βTF1 and βTF2), three Armadillo repeat domains (ARM1, ARM2, and ARM3), a central linker domain (CLD), a juxtamembrane domain (JD), and a short C-terminal domain (CTD)^6-10^ (Fig. 1).

**Fig. 1.**
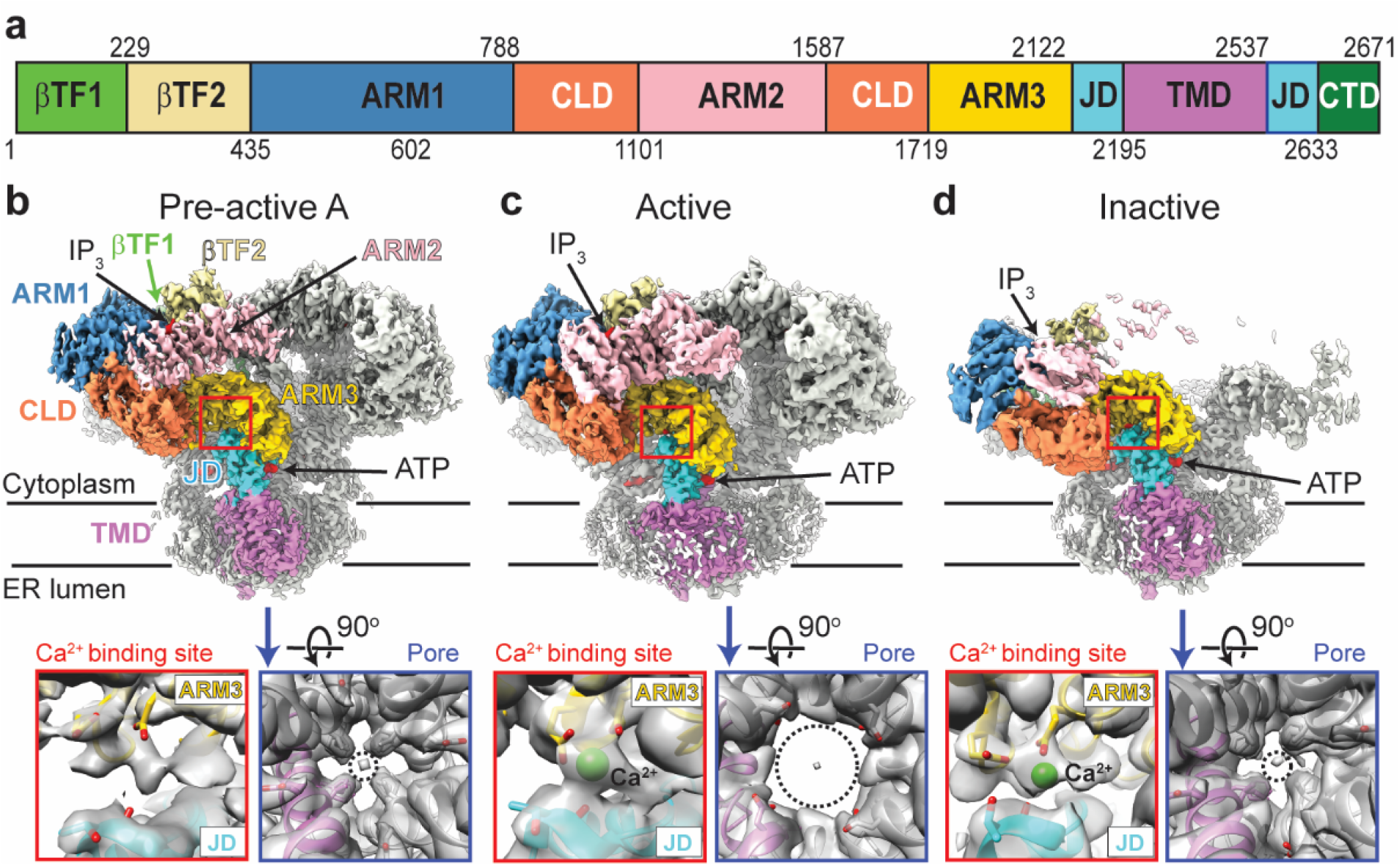
Cryo-EM structures of hIP_3_R-3 in multiple conformations. **a**, Domain boundaries of hIP_3_R-3. **b-d**, Cryo-EM maps of hIP_3_R-3 in pre-active A (**b**), active (**c**), inactive (**d**) conformations. Each domain in one of the subunits is colored as in panel a. Maps within the boxes, shown transparent, are close-up views of the Ca^2+^ binding site (red) and the pore (blue) with ribbon representation of hIP_3_R-3. Select residues are shown in the sticks. Dashed circles indicate opening through the gate.

In addition to IP_3_, the receptor activation requires Ca^2+^ at nanomolar concentrations, whereas Ca^2+^ at higher concentrations is inhibitory, causing the receptor to be tightly regulated by Ca^2+ 11-15^. The cryo-EM structure of human IP_3_R-3 (hIP_3_R-3) in the presence of the inhibitory Ca^2+^ concentrations (2 mM) revealed two binding sites ^6^. However, their role in channel activation and inhibition has remained uncertain. Furthermore, although ATP binding potentiates the receptor by increasing the open probability and duration of the channel openings, the underlying molecular mechanism has not been uncovered^16,17^. In this study, we illuminate the structural framework of receptor activation and channel opening by analyzing five cryo-EM structures of hIP_3_R-3 in the closed-pre-activated, open-activated, and closed-inactivated conformations.

### Cryo-EM structures of hIP_3_R-3 gating conformations

To determine the structure of hIP_3_R-3 in the activated state, we supplemented hIP_3_R-3 with 0.5 mM IP_3_, 5 mM ATP, 1 mM EDTA, and 0.1 mM CaCl_2_ before preparing cryo-grids. Although the free Ca^2+^ concentration was calculated around 100 nM under these conditions^18^, the actual free Ca^2+^ concentration may be higher due to potential leakage of Ca^2+^ during the cryo-grid preparation^6^. We performed a cryo-EM analysis on a large dataset by employing exhaustive 3D classification strategies to separate particles belonging to different functional states resulting in five high resolution (3.2-3.8 Å) structures (Supplementary Fig. 1, 2, Supplementary Table 1). The pore region in all structures resolved to 3.5 Å or better, allowing us to build side chains and determine if the channel was open or closed (Fig. 1, Supplementary Fig. 2). Three structures have closed pores with well-resolved densities for IP_3_ and ATP and are referred to as pre-active A, B, and C (Fig. 1b, Supplementary Fig. 3). The structure named “active” displays drastic conformational changes at the TMD, leading to pore opening (Fig. 1c). In addition to the well-resolved densities for IP_3_ and ATP, the active class reveals substantial density, interpreted as Ca^2+^, at the ARM3-JD Ca^2+^ interface, referred to as the activatory Ca^2+^ binding site (Fig. 1c, Supplementary Fig 3). In the fifth structure, the channel is closed, the activatory Ca^2+^ binding site is occupied, and the intersubunit interactions of the cytoplasmic domains are lost, resembling the hIP_3_R-3 structures obtained in the presence of inhibitory Ca^2+^ concentrations (Fig. 1d)^6^. Therefore, we refer to the structure as “inactive”. Because the map around the second Ca^2+^ binding site identified at high Ca^2+^ concentrations was not resolved well enough to inspect the presence of Ca^2+^, it is unclear if this structure represents a desensitized state that hIP_3_R-3 adopts without additional Ca^2+^ binding or an inhibited state forced by binding of additional Ca^2+^ to an inhibitory site.

### Priming of hIP_3_R-3 for activation

To compare the structures, we aligned their selectivity filters and pore helices (residues 2460-2481), which reside at the luminal side of the TMD and are virtually identical in all classes. Similar to the previous reports^6,19^, IP_3_ binding caused the ARM1 to rotate about 23° relative to the βTF-2, causing global conformational changes within the cytoplasmic domains in the pre-active states compared to the ligand-free structures (Supplementary Fig. 4). Based on the discrete conformational differences observed, a sequential transition from pre-active A to B, then C can be proposed. The pre-active A state is highly similar to the previously published IP_3_-bound hIP_3_R-3 structure^6^. During the transition to the pre-active B state, the N-terminal domain (NTD) of each protomer comprising βTF1, βTF2, ARM1, ARM2, and CLD is rotated about 4° counter-clockwise relative to the TMD and moved about 2 Å closer to the membrane plane (Fig. 2a, Supplementary Video 1). In the pre-active C state, the NTDs remain primarily unchanged, while the ARM3 and JD are rotated by 7°, causing mild distortions at the cytoplasmic side of the TMD without opening the channel (Fig. 2b, Supplementary Video 1). Compared to the ligand-free conformation, the βTFs move about 7 Å closer to the membrane plane, and ARM3-JD rotates about 11° in the pre-active C conformation.

**Fig. 2.**
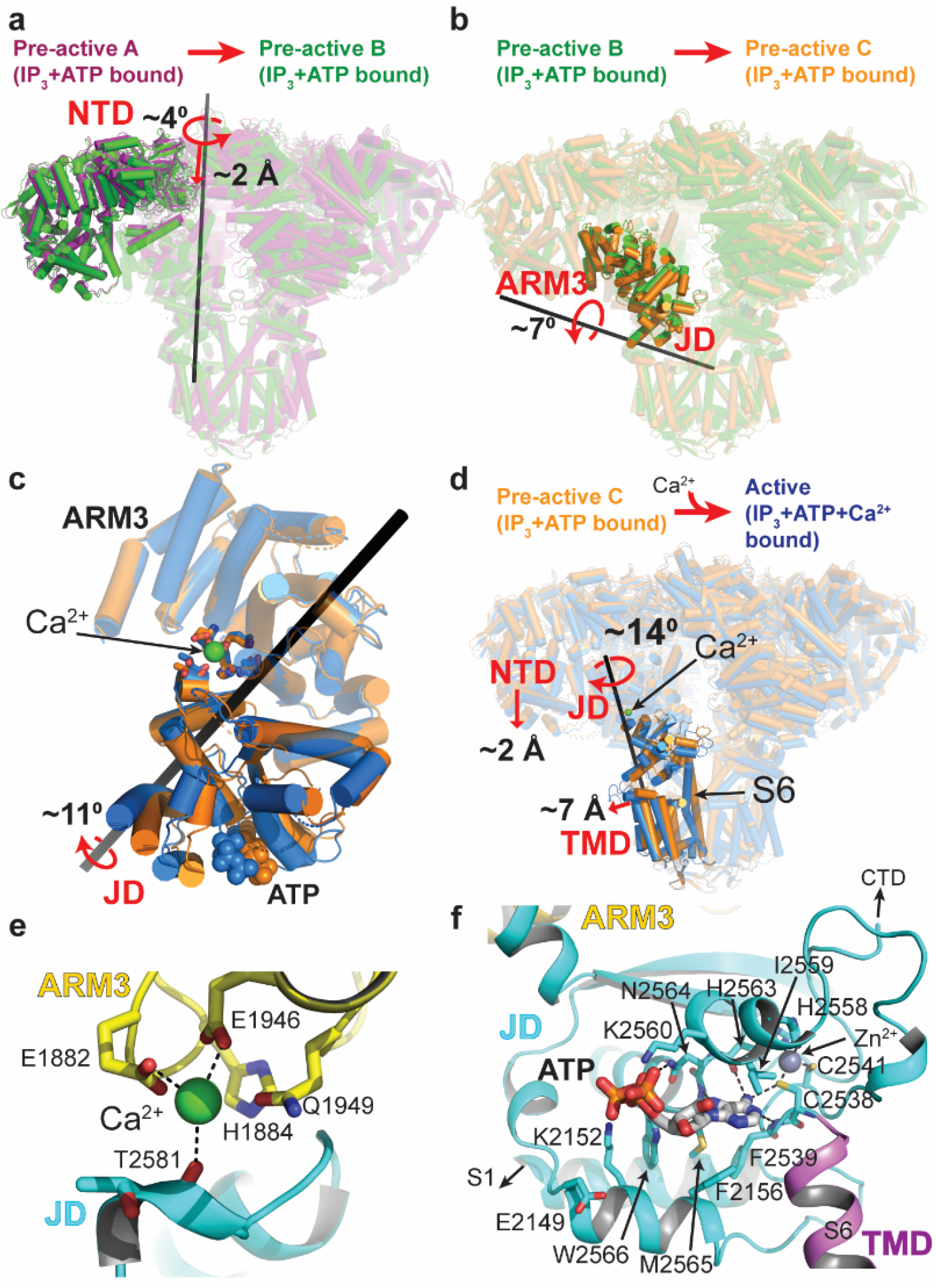
Conformational changes coupling IP_3_ and Ca^2+^ binding to gating. **a**,**b**,**d**, Ribbon representations of hIP_3_R structures superposed on the residues forming the selectivity filter and P-helix of the TMDs, emphasizing the conformational changes between the states indicated above. Domains with substantial conformational changes are shown in full colors only on one subunit, while the rest of the protein is transparent. Curved and straight red arrows indicate the rotation and translation of the domains with red labels relative to the rotation axis (black bars), respectively. **c**, Comparison of the JD (shown in full colors) orientation relative to the ARM3 (shown transparent) in the pre-active-C (orange) and active (blue) structure. The black bar indicates the axis for the rotation of the JD. Ca^2+^ and ATP are shown as spheres. **e**,**f**, Close-up view of the Ca^2+^ binding site in the active conformation and the ATP binding site. Domains are colored as in Fig. 1.

### Ca^2+^-mediated conformational changes leading to channel opening

In the absence of Ca^2+^, the ARM3 and JD act as a rigid body, where there are no significant conformational changes relative to each other (Fig. 2a,b). When Ca^2+^ is bound, the JD rotates about 11°relative to the ARM3 (Fig. 2c), resembling a clamshell closure, which leads to global conformational changes in the whole receptor, including the movement of the NTD closer to the membrane plane by 2 Å (Fig. 2d, Supplementary Video 1). In contrast to the limited rotation of the ARM3 (about 5°), the JD rotates about 14°around an axis roughly perpendicular to the membrane plane, leading to conformational changes at the TMD and resulting in pore opening in the active state (Fig. 2d). Ca^2+^ is coordinated by E1882 and E1946 on the ARM3 and the main-chain carboxyl group of T2581 on the JD (Fig. 2e). H1884 and Q1949 are also close and may interact with Ca^2+^ through water molecules (Fig. 2e). These residues are highly conserved in the homologous ion channel family, ryanodine receptors (RyRs), suggesting a common activation mechanism in IP_3_Rs and RyRs^20^ (Supplementary Fig. 5a,b). Mutation of the corresponding residues in RyRs markedly reduced the sensitivity to Ca^2+^, further supporting this site’s involvement in the Ca^2+^ induced activation^21-23^.

### ATP binding site

Within the JD, we observed a well-resolved cryo-EM density for ATP in all the structures (Fig. 2f, Supplementary Fig. 3). The adenosine base intercalates into a cavity surrounded by F2156, F2539, I2559, M2565, and W2566 near the zinc finger motif and forms hydrogen bonds with the sulfur of C2538, the backbone amide group of F2539, and the carbonyl groups of H2563 and I2559 (Fig. 2f). The phosphate moieties interact with K2152, K2560, and N2564 (Fig. 2f). There are no apparent structural changes around the binding site upon ATP binding, suggesting that ATP’s potentiating effect is likely due to the increased rigidity of the JD (Supplementary Fig. 4a,c). ATP binding site is highly conserved among the subtypes except for E2149 which corresponds to lysine and arginine in IP_3_R-1 and IP_3_R-2 (Supplementary Fig. 4d). A positively charged residue instead of E2149 in the proximity of the phosphate moieties may cause tighter interaction of ATP with IP_3_R-1 and IP_3_R-2, explaining the low binding affinity of ATP to IP_3_R-3 compared to IP_3_R-1 and IP_3_R-2. ATP binds to a similar location near the zinc finger motif in RyRs^20^, but residues forming the binding pocket are distinct (Supplementary Fig. 5a,c).

### Structure of the TMD in the open conformation

The TMD of IP_3_Rs has the same overall architecture of voltage-gated ions channels with a central pore domain, consisting of S5, S6, and pore (P) helix, surrounded by pseudo-voltage-sensor domains (pVSDs), consisting of S1, S2, S3, and S4 helices along with two IP_3_R/RyR specific TM helices (S1’ and S1”) (Fig. 3). In the closed channel, F2513 and I2517 of the S6 helix form two layers of hydrophobic constriction at the pore, blocking the path for the permeation of hydrated ions (Fig. 3). JD’s rotation upon Ca^2+^ binding pushes the pVSD’s cytoplasmic side away from the pore domain by about 7 Å and tilts the cytoplasmic side of the S6 (S6_cyt_) by 12° (Fig. 3a, Supplementary Video 1). Concurrently, the S4-5 linker and S5 helix move away from the S6 helix, thereby inducing a distortion of S6 around the constriction site and moving F2513 and I2517 away from the pore. As a result, the diameter of the water-accessible pore increases to 8 Å, large enough to permeate hydrated cations (Fig. 3b). The flexibility introduced by the neighboring glycine residue (G2514), mutation of which to alanine in IP_3_R-1 is associated with spinocerebellar ataxia 29 (SCA29)^24^, is likely critical to the movement of F2513. The tilting of the S6_cyt_ breaks the salt bridge between D2518 and R2524 of the neighboring subunits, moving D2518 towards the pore while pulling R2524 away, which creates an electronegative path on the cytoplasmic side of the pore (Supplementary Fig. 6). In contrast to the prediction of a π- to α-helix transition at the S6_lum_ during channel opening^6,10^, the π-helix remains intact, and its tip acts as a pivot for the S6_cyt_ tilting and bulging (Fig. 3).

**Fig. 3.**
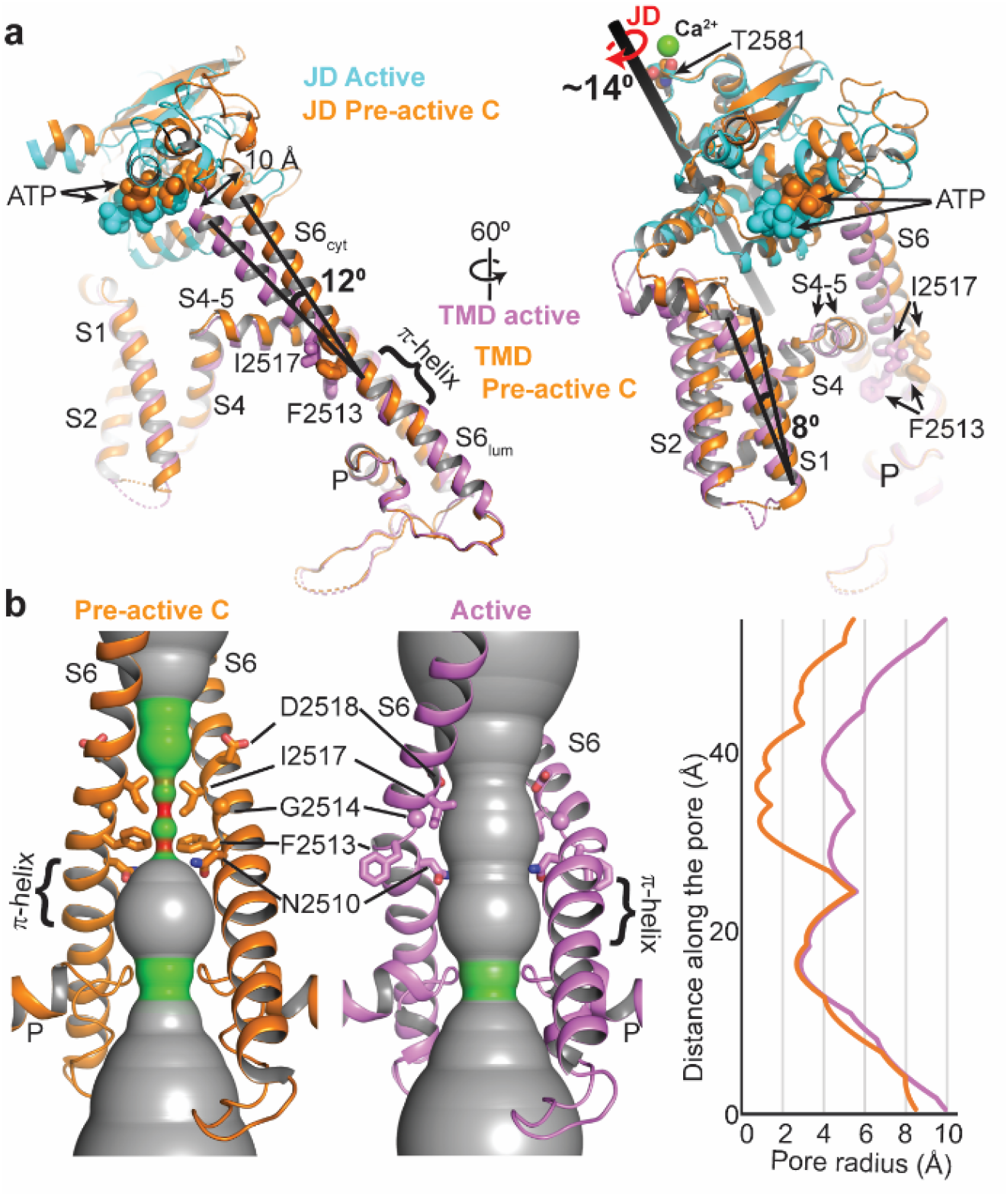
Structure of the IP_3_R-3 in the open conformation. **a**, Comparison of the hIP_3_R-3 structures in the pre-active C and active conformations aligned as Fig. 2. **b**, Ion permeation pathways of hIP_3_R-3 in pre-active C and active conformations (radii coloring: red, <0.8 Å; green, 0.8-4.0 Å; grey, > 4.0 Å) along with the 1D graph of the pore radius. The JD and TMD of the active state are shown in cyan and violet, respectively. The pre-active C state is colored orange.

Although the TMDs of IP_3_Rs and RyRs are highly similar, there are noticeable differences in their pore structures (Supplementary Fig. 5d,e). In RyRs, the constriction site is formed by glutamine and isoleucine, corresponding to F2513 and I2517 in hIP_3_R-3, respectively^25^. In the open state of RyRs, the isoleucine is positioned similarly to I2517 of hIP_3_R-3 ^20,26,27^. On the other hand, the glutamine residue faces the pore in the open state, forming part of the hydrophilic permeation pathway, unlike F2513. Interestingly, N2510 in hIP_3_R-3, which corresponds to alanine in RyRs, faces the permeation pathway similar to the glutamine of RyRs, suggesting that the amide group plays an important role in the ion permeation. However, since the side chain of N2510 extends from a different position on the S6 helix than the side chain of glutamine in RyRs, the binding pocket for ryanodine^20^, a RyR-specific inhibitor, is not present in IP_3_R, potentially causing IP_3_Rs to be unresponsive to ryanodine^25^.

### The flexibility of the CTD

For IP_3_R-1, the CTD was proposed to transmit the conformational changes induced by IP_3_ at the NTD to the JD^8^. However, the CTD is poorly resolved in hIP_3_R-3 structures, including the active state, indicating high flexibility of the CTD relative to the rest of the protein and suggesting that the structural rearrangements of this domain are unlikely to create enough force to open the channel (Supplementary Fig. 7). In line with these observations, removing CTD residues interacting with the βTF2 or swapping the C-terminal region of IP_3_R-1 with the RyRs, which lack the extended CTD, did not diminish receptor activation^19,28,29^.

### Mechanism of hIP_3_R-3 activation and gating

It has been long recognized that IP_3_ binding primes the receptor for activation by Ca^2+ 30^, but how the priming is achieved has remained elusive. Our structures reveal that IP_3_ binding leads to several conformational changes at the NTD, ARM3, and JD, without any apparent structural changes at the activatory Ca^2+^ binding site, and that the ARM3 and JD adopt a new pre-gating conformation relative to the TMD with modest changes of the intersubunit interface between the JDs at the cytoplasmic side of the TMD (Fig. 4, Supplementary Fig. 8, Supplementary Video 1). In addition, ARM3s are constrained in their pre-gating conformation by the tetrameric cage-like assembly of the NTDs, forcing the JDs to rotate upon Ca^2+^ binding. The NTD assembly is maintained by the βTF1-βTF2 intersubunit interactions (βTF ring), which remains intact in the pre-active and active states (Fig. 4, Supplementary Fig. 9, Supplementary Video 1) and acts as a pivot for the conformational changes that stabilize the ARM3. On the other hand, its disruption in the inactive state leads to the loosening of the tetrameric assembly of the NTDs, relieving the ARM3 constraints and causing the JD and TMD to adopt closed channel conformation despite the bound Ca^2+^. Supporting this hypothesis, the removal of βTF1 or mutation of W168, which resides at the βTF1-βTF2 interface (Supplementary Fig. 9), was shown to abolish IP_3_R activity^31,32^.

**Fig. 4.**
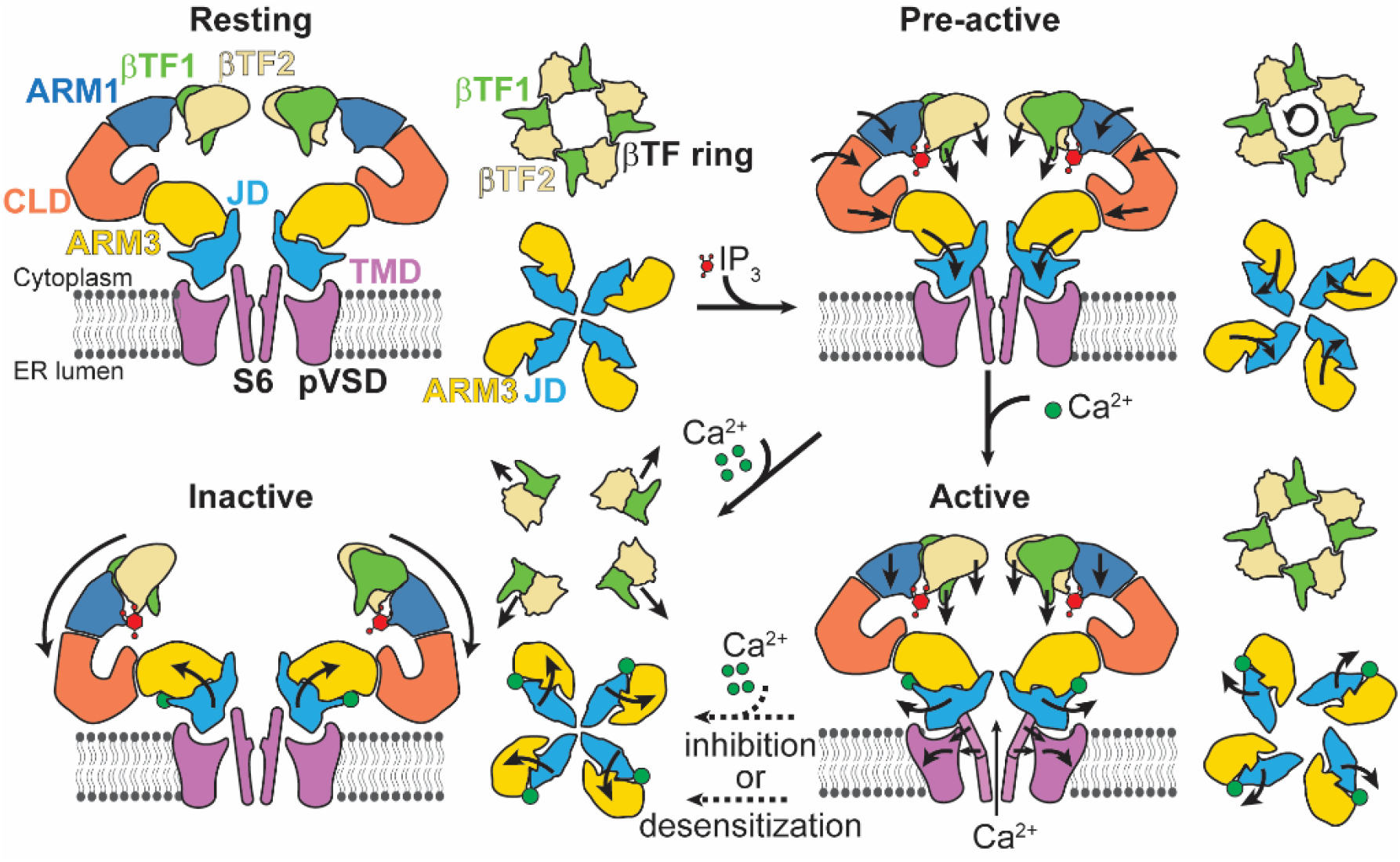
Schematic representation of the IP_3_R gating cycle. A side view of the two opposing subunits (left) and cytoplasmic views of the βTFs and ARM3-JD tetramers (right) for each indicated functional state is depicted. The ARM2 and CTD are omitted for clarity. Arrows shown on the domains indicate the direction of the rotation or translation from the previous conformation.

In conclusion, the ensemble of structures obtained from the same sample demonstrates structural heterogeneity of IP_3_Rs in the presence of IP_3_, ATP, and Ca^2+^. Our ability to correlate these structures with their plausible functional states allowed us to inspect the conformational changes at different gating states, revealing the structural features that define IP_3_R activation and gating. These structures will likely serve as foundations for future experiments addressing biophysical and functional questions related to IP_3_Rs.

## Methods

### Protein expression and purification

Expression and purification of hIP3R-3 were performed as previously described with minor modifications^10^. Briefly, hIP3R-3 (residues 4-2671) with a C-terminal OneStrep tag was expressed using MultiBac expression system. Sf9 cells (4 × 10^6^ cells/mL) were harvested by centrifugation (4,000 g) 48 hours after infection with the baculovirus. Cells resuspended in a lysis buffer of 200 mM NaCl, 40 mM Tris-HCl pH 8.0, 2 mM EDTA pH 8.0, 10 mM β-mercaptoethanol (BME), and 1 mM Phenylmethylsulfonyl fluoride (PMSF) were lysed using Avastin EmulsiFlex-C3. After centrifugation of the lysate at 7,000 g for 20 minutes to remove large debris, the membrane was pelleted by centrifugation at 40,000 rpm (Ti45 rotor) for 1 hour. Membrane pellets were homogenized in ice-cold resuspension buffer (200 mM NaCl, 40 mM Tris-HCl pH 8.0, 2 mM EDTA pH 8.0, 10 mM BME) using a Dounce homogenizer, and solubilized using 0.5% Lauryl maltose neopentyl glycol (LMNG) and 0.1% glyco-diosgenin (GDN) at a membrane concentration of 100 mg/mL. After 4 hours of gentle mixing in the cold room, the insoluble material was pelleted by centrifugation at 40,000 rpm (Ti45 rotor) for 1 hour, and the supernatant was passed through Strep-XT resin (IBA Biotagnology) via gravity flow.

The resin was washed first with 5 column volume (CV) of wash buffer composed of 200 mM NaCl, 20 mM Tris-HCl pH 8.0, 10 mM BME, 0.005% GDN, 0.005% LMNG, followed by 5 CV of wash buffer supplemented with 5 mM ATP and 20 mM MgCl_2_ to remove any bound chaperone proteins, and finally with 5CV of wash buffer supplemented with 1 mM EDTA. The protein was eluted using wash buffer supplemented with 1 mM EDTA and 100 mM D-Biotin (pH 8.2). The protein was further purified by size exclusion chromatography using a Superose 6 column (10/300 GL, GE Healthcare) equilibrated with 200 mM NaCl, 20 mM Tris-HCl pH 8.0, 1 mM EDTA pH 8.0, 2 mM TCEP, 0.005% LMNG, and 0.005% GDN. The fractions corresponding to hIP3R-3 were combined and concentrated to 4 mg/mL using a 100 kDa centrifugal filter (Sartorius). The concentrated sample was then centrifuged at 70,000 rpm using a S100AT rotor (Thermofisher). The concentration dropped to 1.8 mg/mL.

### Cryo-EM sample preparation and data collection

Purified hIP_3_R-3 was supplemented with 500 μM IP_3_ (from 10 mM stock in water), 0.1 mM CaCl_2_, and 5 mM ATP (from 100 mM stock, pH 7.2). 2.0 μL of the protein sample was applied to 300 mesh Cu Quantifoil 1.2/1.3 grids (Quantifoil Microtools) that were glow discharged for 20 seconds at 25 mA. The grids were blotted for 7 s at force 10 using single-layer Whatman filter papers (1442-005, GE Healthcare) before plunging into liquid ethane using an FEI MarkIV Vitrobot at 8 °C and 100% humidity. Four grids prepared using the same sample were imaged using a 300 kV FEI Krios G3i microscope equipped with a Gatan K3 direct electron camera in four different data collection sessions at Case Western Reserve University. Movies containing 40-50 frames were collected at a magnification of 105,000x in super-resolution mode with a physical pixel size of 0.828 Å/pixel and defocus values at a range of −0.8 to −1.6 μm using the automated imaging software SerialEM^33^ and EPU (ThermoFisher Scientific).

### Cryo-EM data processing

Datasets from four sessions were initially processed separately using Relion 3.0 ^34^. We used MotionCor2^35^ and Gtcf^36^ to perform beam-induced motion correction and CTF estimations, respectively. We performed auto picking using the Laplacian-of-Gaussian option of Relion, extracted particles binned 4×4, and performed 2D class classification. Using the class averages with apparent features, we performed another round of particle picking. We cleaned the particles, extracted as 4×4 binned, through 2D classification and performed 3D classification using the hIP_3_R-3 map (EMD-20849^10^), which was converted to the appropriate box and pixel size. We observed two predominant conformations. One had a compact NTD and tight interactions between subunits as in previously published IP_3_R structures in the absence of Ca^2+^, hereafter called “compact” conformation^6-8,10^ (Supplementary Fig. 1). The other one had the NTD of each subunit tilted away from the central symmetry axis resembling hIP_3_R-3 structures obtained in the presence of high Ca^2+^ concentrations, hereafter called “loose” conformation^6^ (Supplementary Fig. 1). These particles were separately selected and reextracted using a box size of 480×480 pixels at the physical pixel size. After 3D refinements, we performed CTF refinement and Bayesian polishing^37^.

We combined all the polished particles and performed another round of 3D classification, using one of the compact structures as a reference map. We grouped particles into “compact” and “loose” classes. Refinement of the particles in the “compact” conformation yielded a 3D reconstruction with an average resolution of 3.9 Å. Although there were slight changes at the TMD compared to the structure in the closed state, these changes were not significant enough to suggest that the channel was open. To more clearly resolve the density around the TMD, we performed another round of 3D classification using a mask that only covers the ARM3, JD, and TMD and without performing an angular or translational alignment in Relion3 ^38^. 3D refinements of the particles in each class were performed using non-uniform refinement in CryoSPARC, enforcing C4 symmetry and local CTF refinements^39^. Four classes led to high-resolution (better than 4Å) 3D reconstructions, whereas the other four classes resulted in poorly resolved maps and were not analyzed further. The particles in the “loose” conformation were processed using non-uniform refinement in CryoSPARC, but non enforcing any symmetry. Local resolution estimates were calculated using CryoSPARC^39^ (Supplementary Fig. 2). Some of the data processing and refinement software was supported by SBGrid^40^.

### Model building

Model building was performed using Coot^41^. We first placed the hIP_3_R-3 structure in ligand-free conformations (PDB ID:6UQK^10^) into the density map of Pre-active A, and performed rigid-body fitting of individual domains of one of the protomers. We then manually fit the residues into the density and expand the protomer structure into a tetramer using the C4 symmetry. We performed real-space refinement using Phenix^42^. We repeated build-refine iterations till a satisfactory model was obtained. This model was used as a starting model for the other structures following the same workflow. Regions without interpretable densities were not built into the model. Residues without apparent density for their side chains were built without their side chains (*i*.*e*., as alanines) while maintaining their correct labeling for the amino acid type.

### Figure preparation

Figures were prepared using Chimera^43^, ChimeraX^44^, and The PyMOL Molecular Graphics System (Version 2.0, Schrödinger, LLC). Calculation of the pore radii was performed using the software HOLE^45^.

## Acknowledgments

We thank Dr. Kunpeng Lee for cryo-EM data collection at Case Western Reserve University. We thank Theo Humphries and other support staff at the Pacific Northwest Center for Cryo-EM (PNCC), Drs. Elad Binshtein, Melissa Chambers, and Scott Collier at the Cryo-EM facility at Vanderbilt University for their assistance with cryo-EM sample screening. We thank Drs. Hassane Mchaourab, Terunaga Nakagawa, and Silvia Ravera for discussions and review of the manuscript. This work was conducted in part using the CPU and GPU resources of the Advanced Computing Center for Research and Education (ACCRE) at Vanderbilt University. We used the DORS storage system supported by the NIH (S10RR031634 to Jarrod Smith). E.K. was supported by Start-up fund from Vanderbilt University and Vanderbilt Diabetes and Research Training Center (DK020593). T.N. was supported by the NIH (R01HD061543) and the Vanderbilt University. C.M.A. and E.A.L. were supported by the Molecular Biophysics Training Program (T32 GM008320 to Walter Chazin).

## Author contributions

E.K. conceived the project and performed cryo-EM data analysis; EAS optimized and performed protein expression and purification; HT performed grid preparation for cryo-EM. All authors contributed to the preparation of the manuscript.

## Competing interests

The authors declare no competing interests.

## Materials and Correspondence

Reagents and other materials will be available upon request from EK with a completed materials transfer agreement.

**Supplementary Fig. 1.**
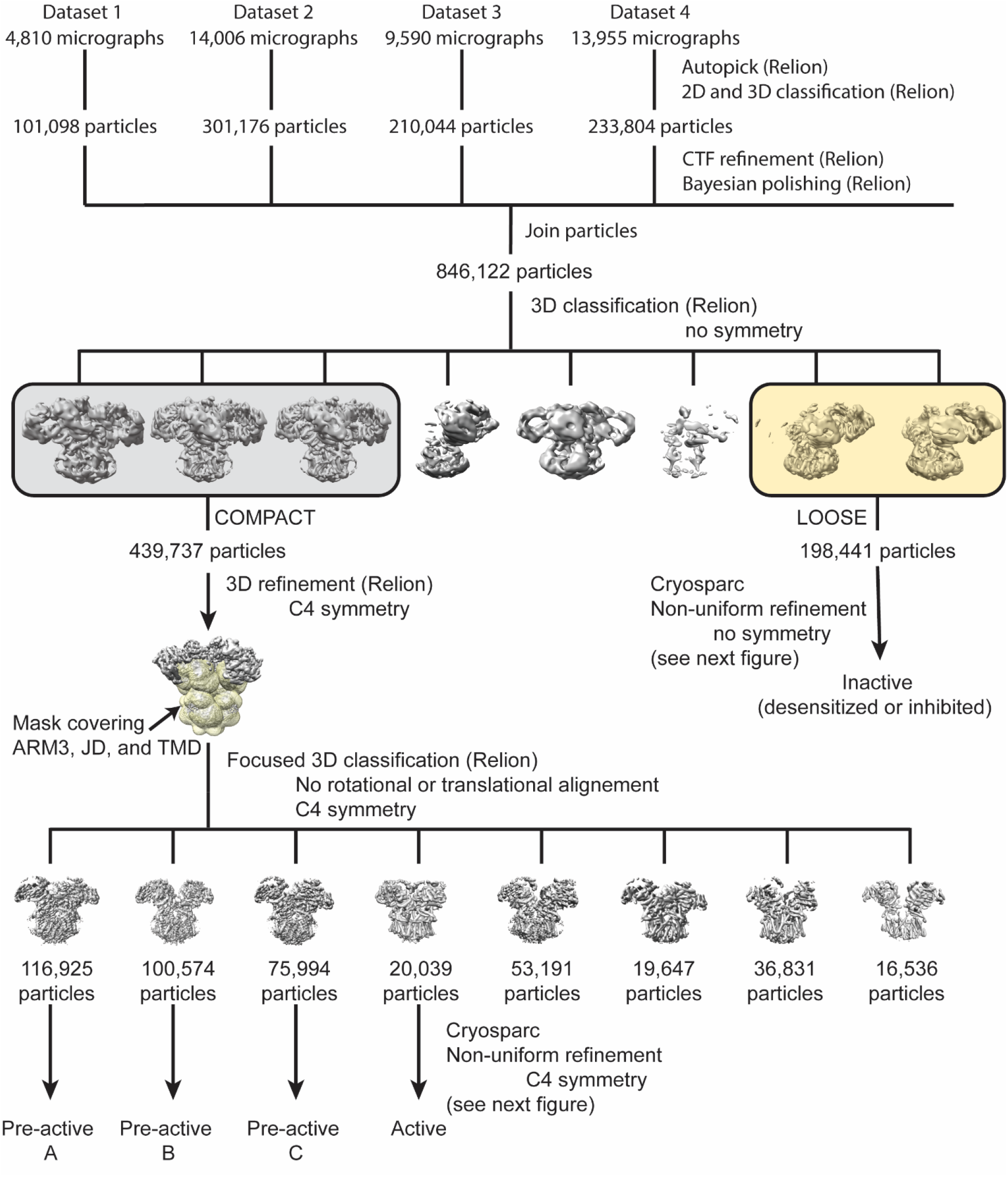
Cryo-EM analysis of hIP3R-3. Flowchart detailing the particle selection and refinement procedure to obtain the cryo-EM maps of hIP_3_R-3 in different gating conformations. See methods for details.

**Supplementary Fig. 2.**
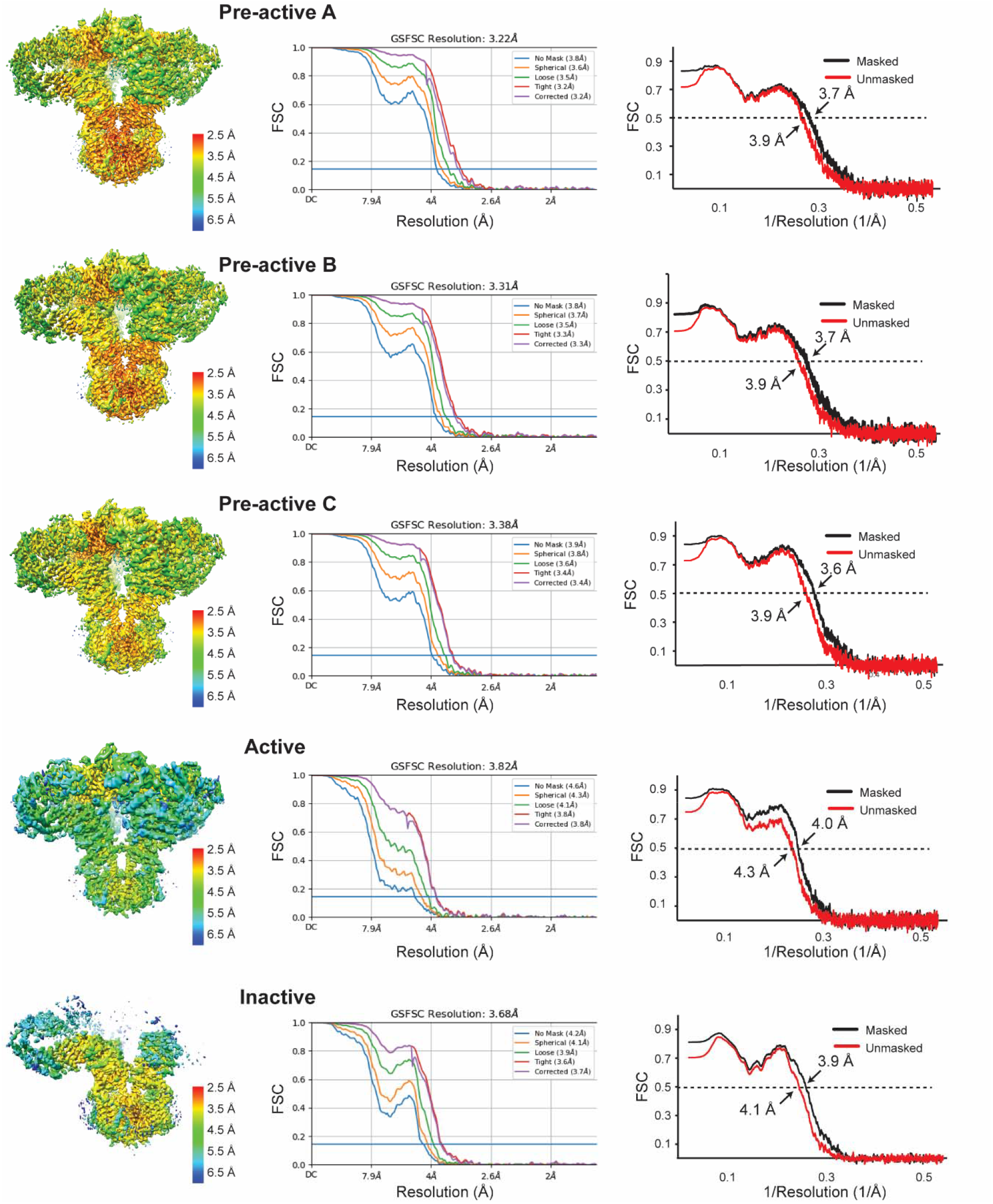
Cryo-EM analysis of hIP3R-3. Local resolution mapped onto the final refined 3D reconstructions of hIP3R-3 are shown on the left for each structure. FSC curves after Non-uniform refinement in CryoSPARC are shown in the middle, and FSC curves of the refined model versus the EM map are shown on the right.

**Supplementary Fig. 3.**
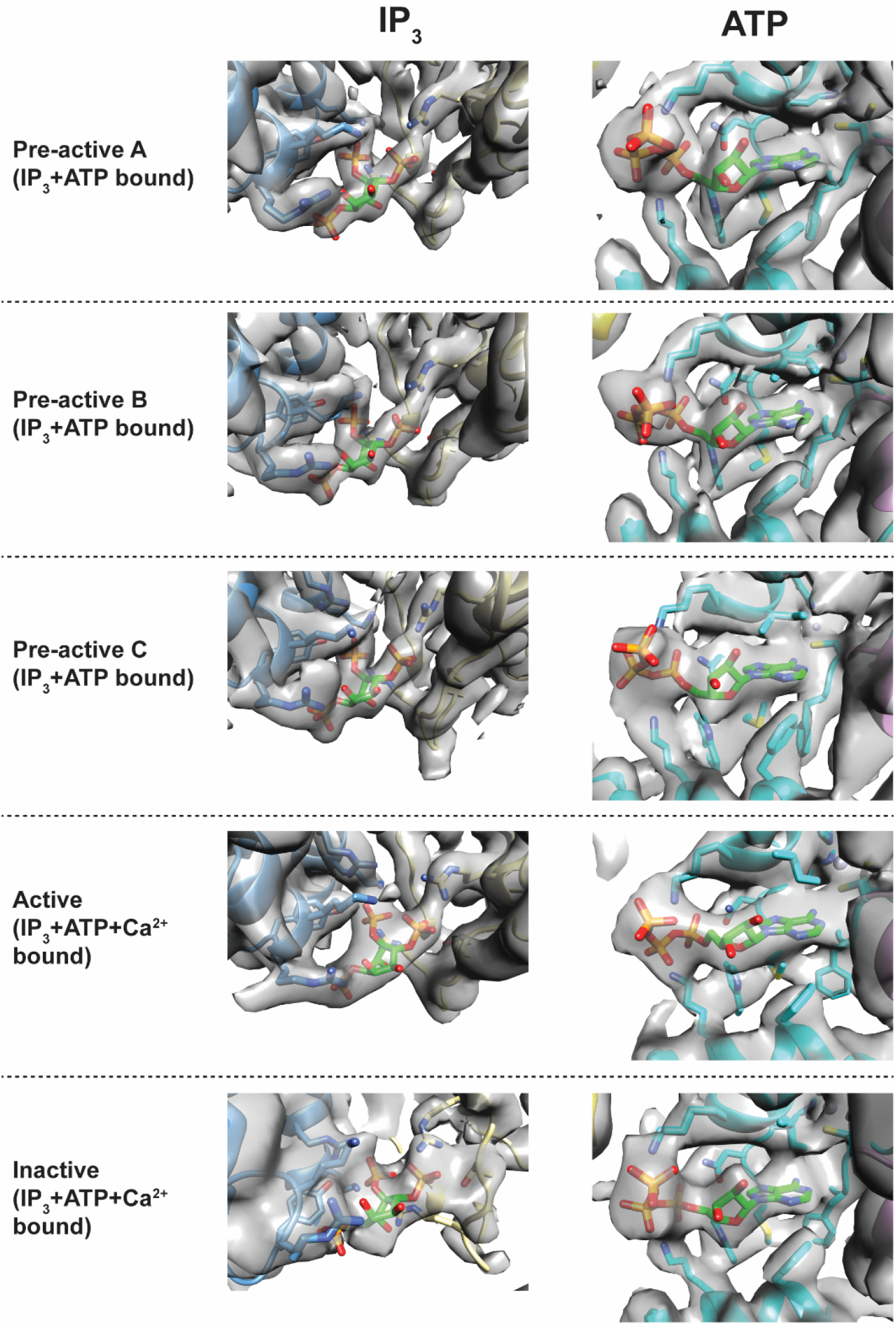
Ligand binding sites. Density maps for the IP_3_ and ATP binding site are shown as transparent grey surfaces. The ligands and the neighboring residues are represented as sticks.

**Supplementary Fig. 4.**
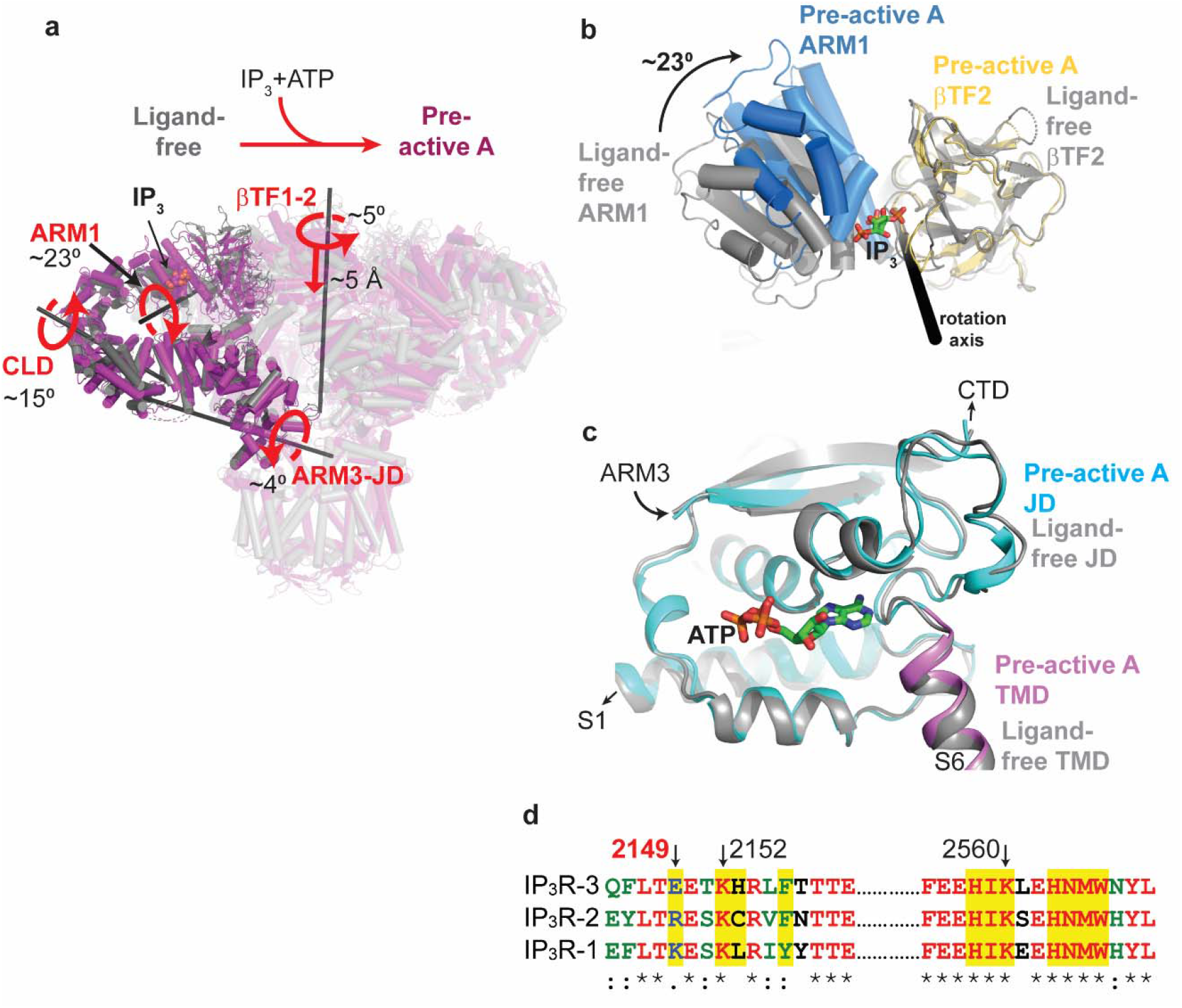
Structural comparison of the hIP_3_R-3 structures in the ligand-free and Pre-active A conformations. **a**, Ribbon representations of hIP_3_R structures in the resting (PDB ID: 6UQK^10^) superposed on the residues forming the selectivity filter and P-helix of the TMDs, emphasizing the conformational changes upon IP_3_ and ATP binding. Domains with substantial conformational changes are shown in full colors only on one subunit, while the rest of the protein is transparent. Curved and straight red arrows indicate the rotation and translation of the domains with red labels relative to the rotation axis (black bars)., respectively. **b**, IP_3_ induced conformational changes resulting in 23°rotation of the ARM1 relative to βTF2. The structures are aligned on the βTF2. Pre-active A structure is colored as in Fig.1, and Ligand-free structure is colored in grey. IP_3_ is shown as green sticks. **c**, Close-up view of the ATP binding site. Pre-active A structure is colored as in Fig.1, and Ligand-free structure is colored in grey. ATP is shown as green sticks. No major conformational changes around the binding site are observed. **d**, Sequence alignment of hIP_3_R subtypes around the residues forming the ATP binding site. Residues shown in Fig. 2 are highlighted. Select residues are indicated by arrows, and E2149 is labeled in red.

**Supplementary Fig. 5.**
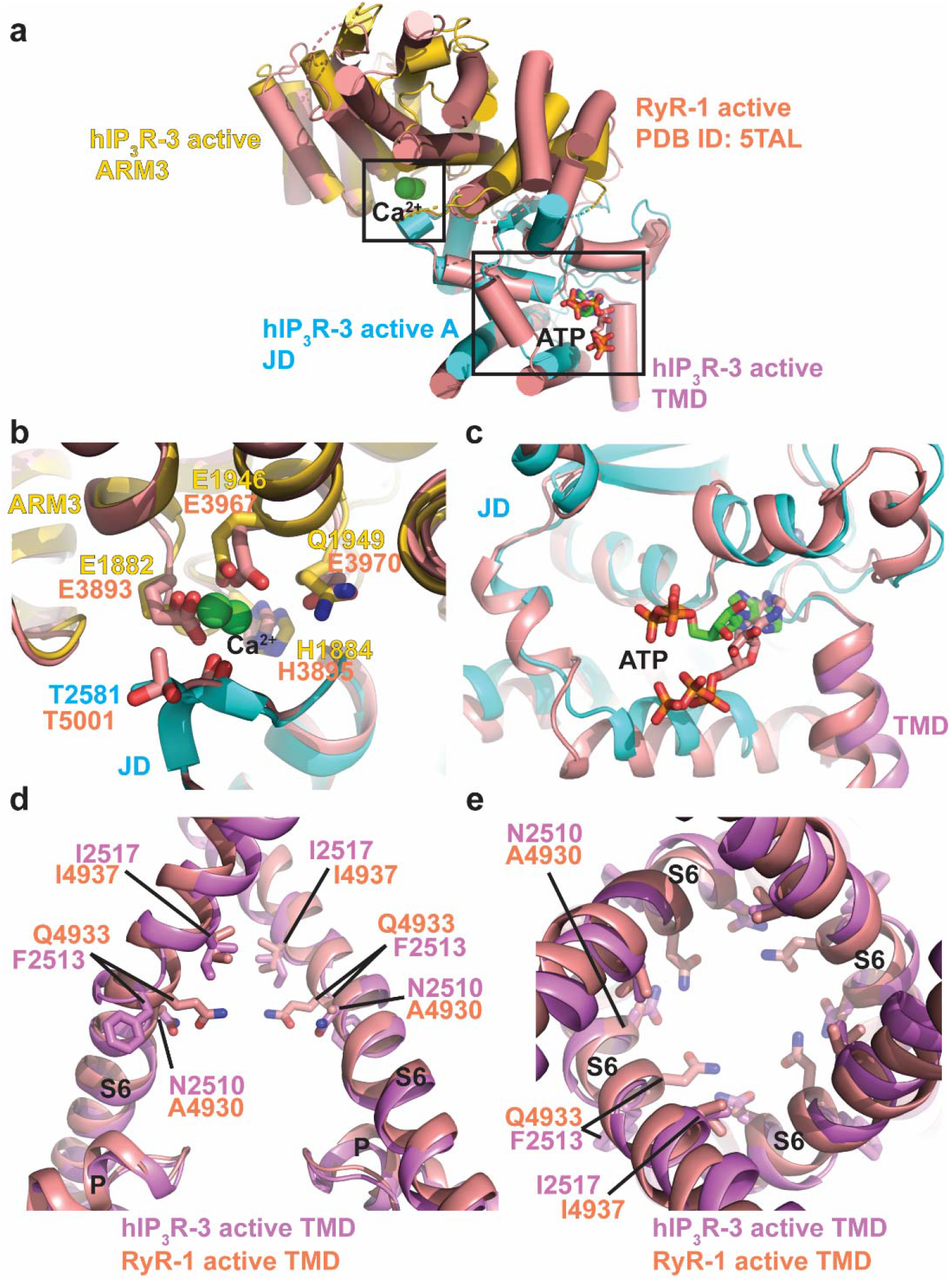
Structural comparison of hIP_3_R-3 and RyR1 structures in the active state. **a**, An overlay of the ARM3, JD, and part of S6 of hIP_3_R-3 with the corresponding domains of rabbit RyR-1 (PDB ID: 5TAL) ^20^. hIP_3_R-3 is colored as in Fig. 1, and RyR1 is colored in salmon. Ca^2+^ and ATP binding sites are boxed. **b-c**, Close-up views of the Ca^2+^ (**b**) and ATP (**b**) binding sites. Residues coordinating Ca^2+^ are shown as sticks and labeled. **d-e**, Alignment of the S6 and P helices viewed through the membrane plane (**d**) and cytoplasm (**e**). Two of the subunits were not shown in panel **d** for clarity.

**Supplementary Fig. 6.**
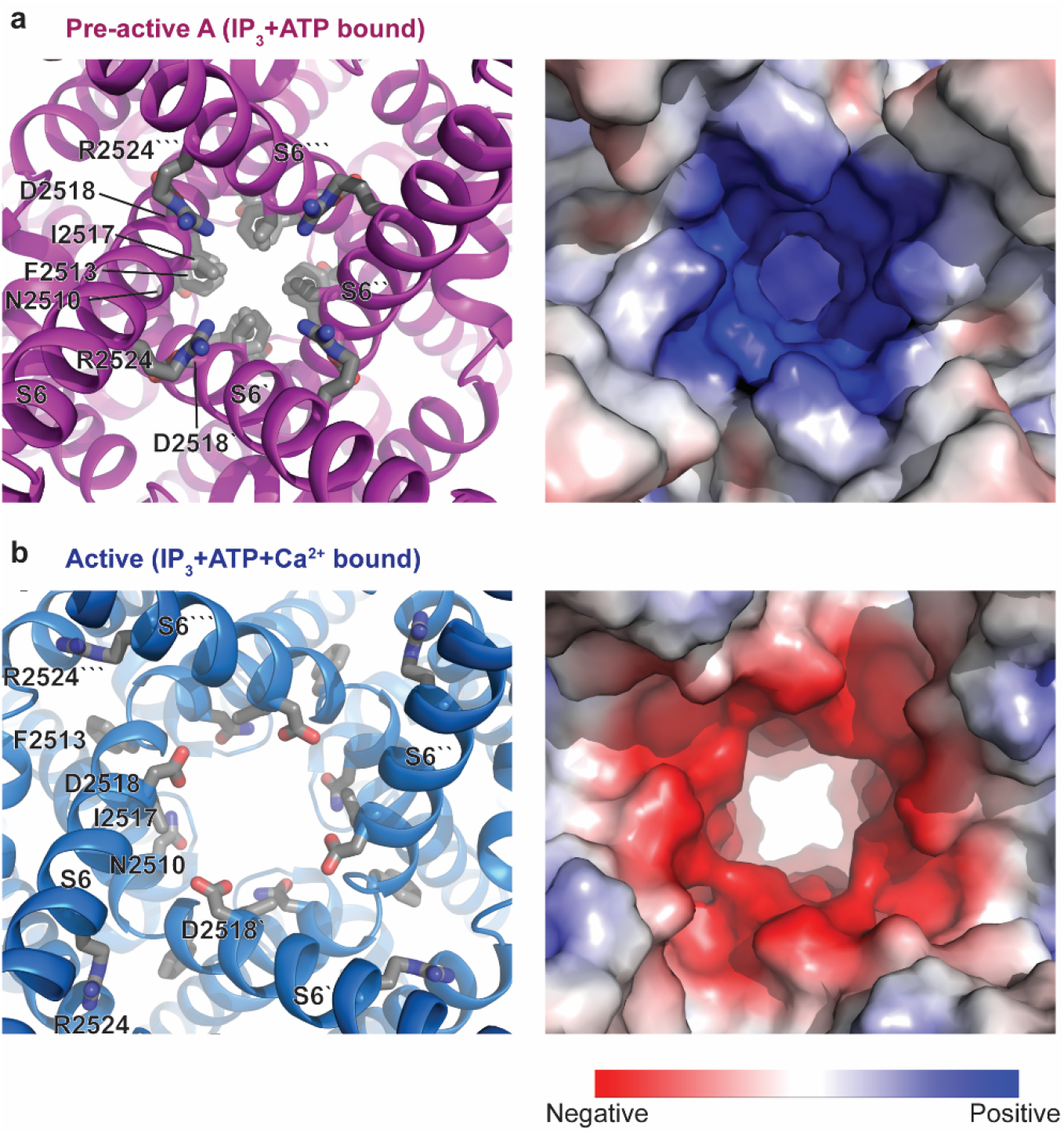
Structural comparison of hIP_3_R-3 pores in closed and open conformations. **a-b**, Close up cytoplasmic view of the IP_3_R-3 pores in the closed (**a**) and open (**b**) conformations in ribbon (left) and electrostatic surface representations. Pore lining residues are shown as grey sticks. ‘is used the differentiate residues in different subunits.

**Extended Fig. 7.**
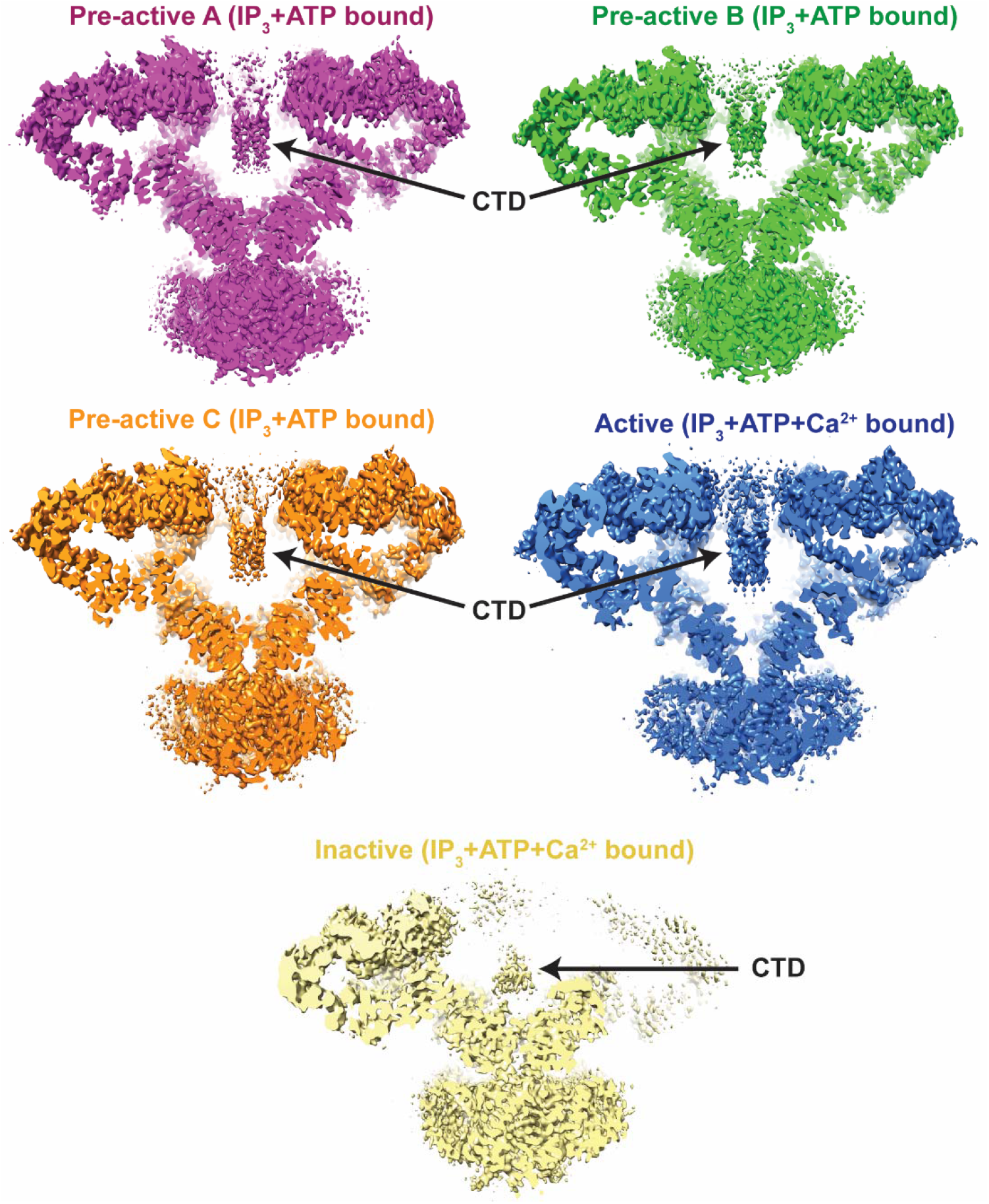
The flexible architecture of the CTD. Cryo-EM map sections of hIP_3_R-3 in different gating conformations focusing on the density for the CTD.

**Supplementary Fig. 8.**
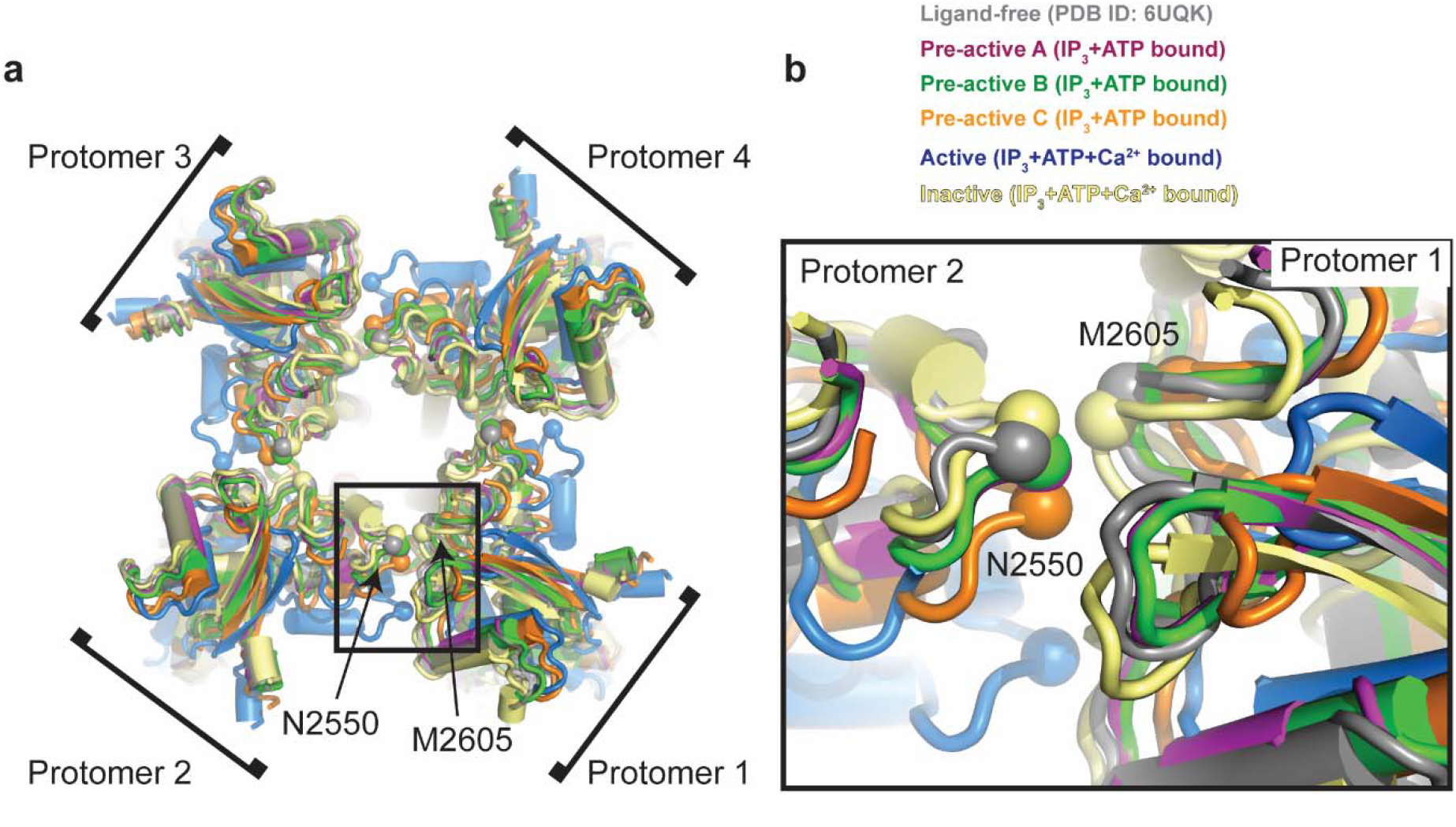
Structural changes of the JDs. **a**, The tetrameric JDs of hIP_3_R-3 in multiple gating conformations viewed from cytoplasm through the 4-fold symmetry axis. Structures are superposed on the residues forming the selectivity filter and P-helix of the TMDs. **b**, Zoomed view of the boxed area. Structures are colored as in Fig. 2. The Cα atoms for the residues N2550 and M2605 are shown as spheres to emphasize the structural differences among different conformations.

**Supplementary Fig. 9.**
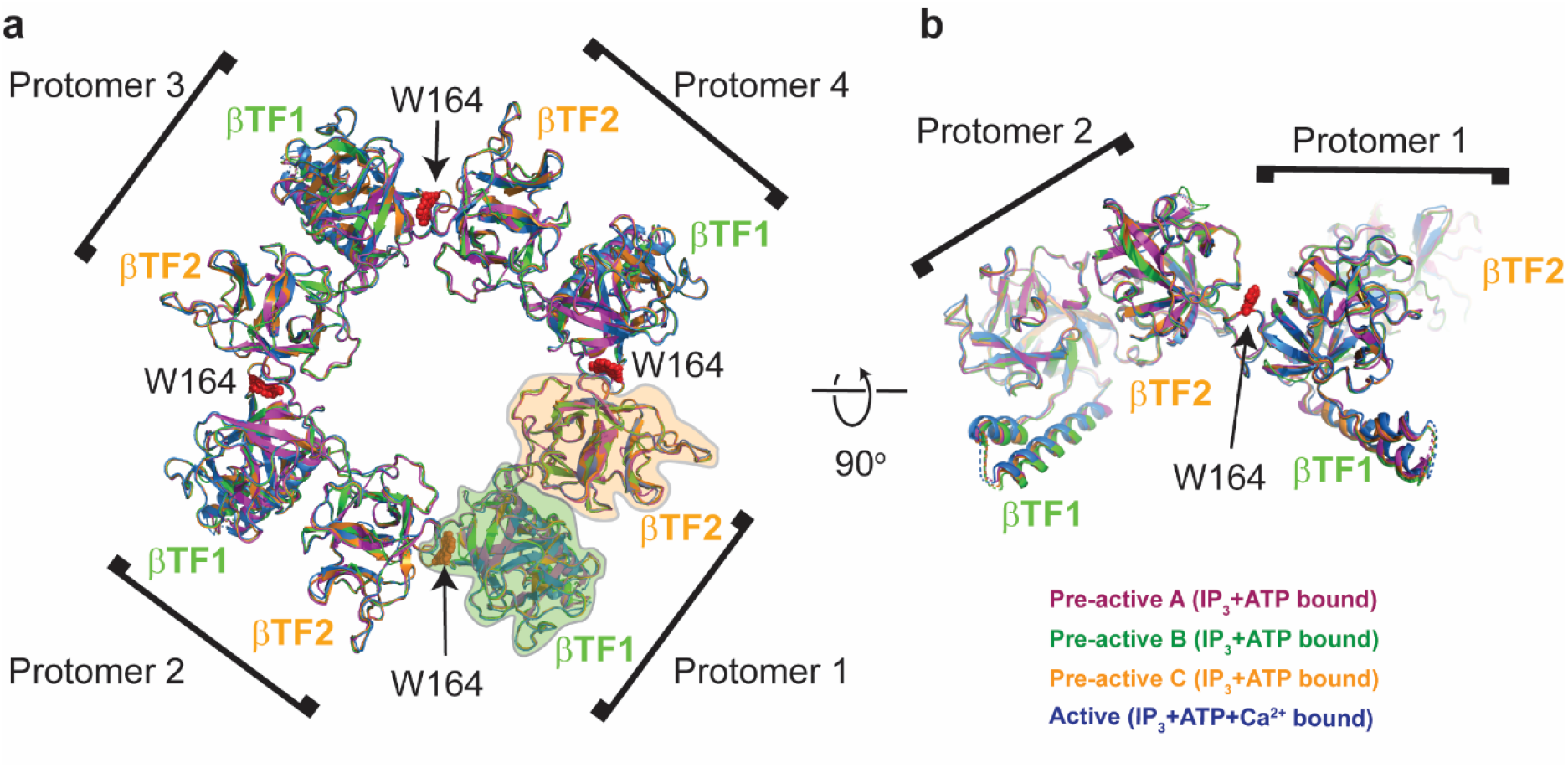
Structure of the βTF ring. **a**, Aligned structures of the βTF rings of hIP_3_R-3 in multiple gating conformations viewed from cytoplasm through the 4-fold symmetry axis. **b**, 90°rotated view of panel **a**. Structures are colored as in Fig. 2. The βTF1 and βTF2 of each protomer are labeled, and they are circled for one of the protomers to highlight their boundaries. W164 is shown as red spheres.

**Supplementary Table 1.** Cryo-EM data collection, refinement, and validation statistics.

| **Data collection and processing** |  |  |  |  |  |
| --- | --- | --- | --- | --- | --- |
| Microscope | FEI Krios G3i microscope |  |  |  |  |
| Detector | Gatan K3 direct electron camera |  |  |  |  |
| Nominal magnification | 105,000 x |  |  |  |  |
| Voltage (kV) | 300 |  |  |  |  |
| Electron exposure (e/Å<sup>2</sup>) | ~60 |  |  |  |  |
| Defocus range (μm) | -0.8 to -1.6 |  |  |  |  |
| Pixel size (Å) | 0.828 |  |  |  |  |
| Number of Micrographs | 42,361 |  |  |  |  |
| Particles images (no.) | 846,122 |  |  |  |  |
| Conformational state | **Pre-active A** | **Pre-active B** | **Pre-active C** | **Active** | **Inactive** |
| Symmetry imposed | C4 | C4 | C4 | C4 | C1 |
| Final particles images (no.) | 116,925 | 100,754 | 75,994 | 20,039 | 198,441 |
| Map resolution (Å) (FSC threshold=0.143) | 3.2 | 3.3 | 3.4 | 3.8 | 3.7 |
| **Refinement** |  |  |  |  |  |
| Model resolution(Å) (FSC threshold=0.5) | 3.7 | 3.7 | 3.6 | 4.0 | 3.9 |
| B-factor used for map sharpening (Å<sup>2</sup>) | -100 | -100 | -100 | -100 | -100 |
| **Model composition** |  |  |  |  |  |
| Non-hydrogen atoms | 55,340 | 56,720 | 60,104 | 56,388 | 24,487 |
| Protein residues | 7,524 | 7,540 | 7,744 | 7,720 | 3545 |
| Zn<sup>2+</sup> | 4 | 4 | 4 | 4 | 4 |
| IP<sub>3</sub> | 4 | 4 | 4 | 4 | 1 |
| ATP | 4 | 4 | 4 | 4 | 4 |
| Ca<sup>2+</sup> | - | - | - | 4 | 3 |
| **Mean B factors (Å<sup>2</sup>)** |  |  |  |  |  |
| Protein | 27.11 | 23.63 | 59.69 | 109.96 | 54.99 |
| Ligands | 20.95 | 19.04 | 44.71 | 104.12 | 72.32 |
| **R.m.s. deviations** |  |  |  |  |  |
| Bond lengths (Å) | 0.002 | 0.002 | 0.003 | 0.002 | 0.004 |
| Bond angles (°) | 0.472 | 0.464 | 0.526 | 0.452 | 0.624 |
| **Molprobity score** | 1.58 | 1.64 | 1.65 | 1.54 | 1.75 |
| **Clash score** | 6.04 | 6.79 | 7.70 | 5.26 | 8.5 |
| **Poor rotamers (%)** | 0.46 | 0.15 | 0.45 | 0.31 | 0.00 |
| **Ramachandran plot** |  |  |  |  |  |
| Favored (%) | 96.32 | 96.12 | 96.54 | 96.22 | 95.75 |
| Allowed (%) | 3.68 | 3.88 | 3.46 | 3.78 | 4.25 |
| Disallowed (%) | 0.00 | 0.00 | 0.00 | 0.00 | 0.00 |

**Supplementary Video 1. Conformational changes during hIP**_**3**_**R-3 activation and gating**.

This video shows a morph of the hIP_3_R-3 structure from ‘pre-active A’ to ‘pre-active B’, ‘pre-active B’ to ‘pre-active C’, and ‘pre-active C’ to ‘active’ states viewed from the side (left) and top (right). Two opposing subunits are colored as individual domains, similar to Fig. 1. The other two subunits are colored in grey. IP_3_ and ATP are shown as spheres and colored red. The gate-forming residues F2513 and I2517 are shown as sticks. Brackets indicate the depth of the view for the movies on the right. A green sphere is used to indicate the Ca^2+^ binding site.

## Notes

### Competing Interest Statement

The authors have declared no competing interest.

